# CRISPR-Cas9 screens in human cells and primary neurons identify modifiers of *C9orf72* dipeptide repeat protein toxicity

**DOI:** 10.1101/129254

**Authors:** Michael S. Haney, Nicholas J. Kramer, David W. Morgens, Ana Jovičić, Julien Couthouis, Amy Li, James Ousey, Rosanna Ma, Gregor Bieri, Michael C. Bassik, Aaron D. Gitler

## Abstract

Hexanucleotide repeat expansions in the *C9orf72* gene are the most common cause of amyotrophic lateral sclerosis and frontotemporal dementia (c9FTD/ALS). The nucleotide repeat expansions are translated into dipeptide repeat (DPR) proteins, which are aggregation-prone and may contribute to neurodegeneration. Studies in model organisms, including yeast and flies have converged upon nucleocytoplasmic transport as one underlying pathogenic mechanism, but a comprehensive understanding of the molecular and cellular underpinnings of DPR toxicity in human cells is still lacking. We used the bacteria-derived clustered regularly interspaced short palindromic repeats (CRISPR)-Cas9 system to perform genome-wide gene knockout screens for suppressors and enhancers of *C9orf72* DPR toxicity in human cells. We validated hits by performing secondary CRISPR-Cas9 screens in primary mouse neurons. Our screens revealed genes involved in nucleocytoplasmic transport, reinforcing the previous findings from model systems. We also uncovered new potent modifiers of DPR toxicity whose gene products function in the endoplasmic reticulum (ER), proteasome, RNA processing pathways, and in chromatin modification. Since regulators of ER stress emerged prominently from the screens, we further investigated one such modifier, *TMX2*, which we identified as a modulator of the ER-stress signature elicited by *C9orf72* DPRs in neurons. Together, this work identifies novel suppressors of DPR toxicity that represent potential therapeutic targets and demonstrates the promise of CRISPR-Cas9 screens to define mechanisms of neurodegenerative diseases.

**One Sentence Summary:** Genome-wide CRISPR-Cas9 screens in human cells reveal mechanisms and targets for ALS-associated *C9orf72* dipeptide repeat protein toxicity.

---

The ALS and FTD causing mutation in the *C9orf72* gene is a massively expanded hexanucleotide repeat (GGGGCC) (*1, 2*), which produces sense and antisense RNA foci (*1*) and is translated into aggregation-prone dipeptide repeat (DPR) proteins via an unconventional form of AUG-independent translation (also called RAN translation) (*3-5*). Studies in flies and human cells suggest DPRs may be the main drivers of neuronal toxicity (*6-8*). The arginine-rich DPRs, Glycine-Arginine (GR) and Proline-Arginine (PR) are particularly toxic in experimental models (*6, 8-11*). Synthetic PR or GR DPRs added exogenously to the culture media are rapidly transported to the nucleus, cause disruptions in RNA splicing - including in the canonical splicing and biogenesis of ribosomal RNA (rRNA) - and induce cell death in a dose-dependent manner (*9*). Subsequent studies have provided evidence through co-immunoprecipitation and mass-spectroscopy that these DPRs preferentially bind proteins with low complexity domains, including RNA-binding proteins (*12, 13*), ribosomal proteins, and translational elongation factors (*14, 15*), as well as nuclear pore complex components (*16*).

Genetic screens in simple experimental model organisms like yeast, flies, and worms have empowered the discovery of fundamental biological processes including mechanisms of human disease (*17*). Recent technological advances in CRISPR-Cas9 genome editing have expanded the scope and reliability of genome-wide genetic deletion screens to the human genome using high complexity single-guide RNA (sgRNA) libraries (*18-21*). Here, we used the CRISPR-Cas9 system to perform comprehensive genome-wide knockout screens in human cells and mouse primary neurons to identify genetic modifiers of *C9orf72* DPR toxicity.

We engineered the human immortalized myelogenous leukemia cell line K562 to stably express Cas9 (*22*). Similar to previous reports in other human cell lines (*9*), synthetic polymers of PR and GR (PR_20_ and GR_20_) added to cell culture media were rapidly taken up by K562 cells, trafficked to the nucleus, and killed cells in a dose-dependent manner (**Fig. 1A-D, Fig. S1**). To identify genetic modifiers of both PR_20_ and GR_20_ mediated-toxicity, we conducted genome-wide CRISPR knockout screens. We used a lentiviral sgRNA library comprised of ten sgRNAs per gene, targeting ∼20,500 human genes, along with ∼10,000 negative control sgRNAs (*23*) (**Fig. 1E**). We used deep sequencing to track the effect (protective, sensitizing, neutral) of each sgRNA in a pooled population of knockout cells. That is, sgRNAs that protected cells from DPR toxicity were enriched and those that sensitized cells were depleted from the pool. In order to identify both protective and sensitizing gene knockouts in the same screen, we treated the pooled population of cells with a dose of DPRs sufficient to kill 50% of cells and repeated this treatment four times over the course of the screen to amplify the selection. We performed separate screens for modifiers of PR_20_ and GR_20_ toxicity, each in duplicate, and repeated each screen two independent times.

**Figure 1.**
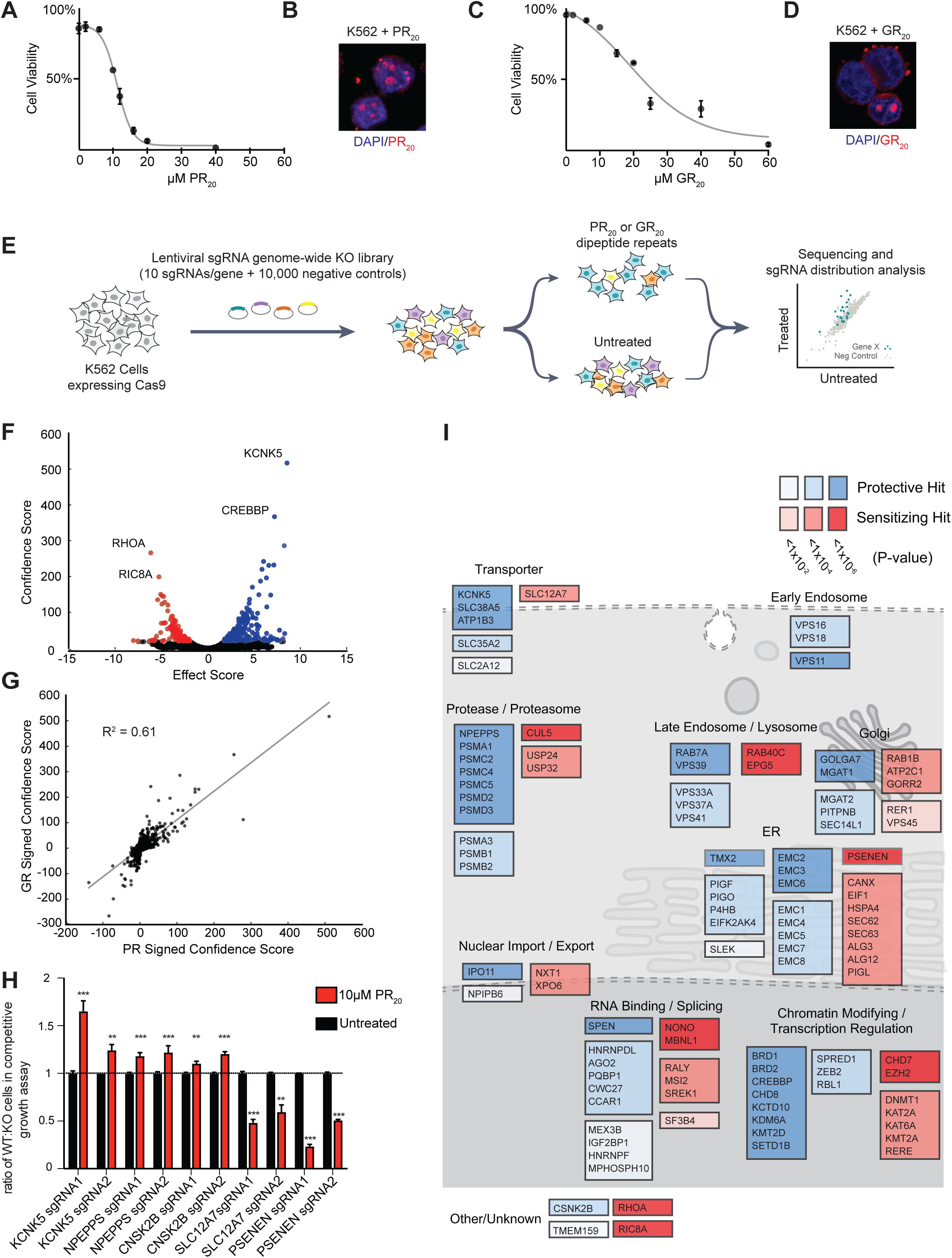
Genome-wide CRISPR-Cas9 knockout screens in human cells identify modifiers of *C9orf72* DPR toxicity. FLAG-tagged synthetic DPRs (PR_20_ or GR_20_) were added to the culture media of K562 cells, which resulted in dose dependent cytotoxicity (**A, C**). The DPRs were internalized and localized to the nucleus of cells as visualized by immunocytochemistry (**B, D,** blue = DAPI, red = anti-FLAG). **(E)** Pooled CRISPR-Cas9 screening paradigm. K562 cells stably expressing Cas9 were infected with a lentiviral sgRNA library, the population was split and 1/2 was treated with PR_20_ or GR_20_ for 4 pulses, and then the resulting populations were subjected to deep sequencing and analysis. **(F)** Volcano plot for all human genes in the GR_20_ screen. Colored in blue are all the genes conferring resistance to GR_20_ when knocked out (10% FDR) and colored in red are all the genes conferring sensitivity to GR_20_ when knocked out (10% FDR). **(G)** Correlation of signed confidence scores of significant hits between GR_20_ and PR_20_ screens (R^2^ = 0.61). **(H)** Validation of hits using independently generated single gene knockout K562 lines in co-culture competitive growth assays. Equal numbers of knockout cells (GFP^+^) and WT control cells (GFP–) were co-cultured and subsequently treated with PR_20_ (10uM). The abundance of knockout cells (GFP^+^ cells) was quantified by flow-cytometry 48 hours after PR_20_ treatment and compared to the untreated population (two-tailed t-test; *** p < 0.001, ** p < 0.01) Each gene was validated using two distinct sgRNAs. **(I)** Schematic of selected hits (10% FDR) from both PR_20_ and GR_20_ screens colored coded by significance (p-value) and depicted at subcellular localizations based on gene ontology.

We then used the casTLE algorithm as previously described (*23*) to detect statistically significant suppressors and enhancers of DPR toxicity. Using a false discovery rate (FDR) cut-off of 10%, the screens identified 215 genetic modifiers of PR_20_ toxicity and 387 genetic modifiers of GR_20_ toxicity (**Fig. 1F, S2A-C, Supplemental Table 1**). Results from the PR_20_ and the GR_20_ screen were well correlated (R^2^=0.61), suggesting similar mechanisms of toxicity (**Fig. 1G**). We validated individual knockout K562 lines by performing competitive growth assays with PR_20_ treatment (**Fig. 1H**). Among the hits from these knockout screens were genes encoding some of the same nuclear import and export factors discovered in DPR modifier screens in model organisms (*10, 11, 24, 25*), such as *XPO6* and *IPO11* (**Fig. 1I**), suggesting the same mechanisms are important in mammalian cells. In addition, we identified many novel genetic modifiers of PR and GR toxicity, including genes encoding RNA-binding proteins that have been shown to physically interact with PR and GR (*12*), such as *NONO* and *HNRNPF*. We also identified genes involved in RNA-splicing such as *SPEN* and *PQBP1*, a process previously observed to be dysregulated in C9FTD/ALS patient brains (*26*) and dramatically disrupted in human cell lines treated with DPRs (*9*). In addition to these previously implicated genes, some of the strongest hits in the screen were genes encoding ER-resident proteins, such as *TMX2, CANX*, almost all members of the endoplasmic reticulum membrane protein complex (EMC), a complex that is upregulated by ER stress and is thought to be involved in ER-associated degradation (ERAD) (*27-29*), and genes encoding proteasome subunits. Chromatin modifiers and transcriptional regulators, including genes involved in histone lysine methylation (*KDM6A*, *KMT2D*, *KMT2A*, and *SETD1B*) were also significant genetic modifiers of PR and GR toxicity.

To evaluate the role of the genetic modifiers identified in our human genome-wide screens in a more disease relevant context, we next designed a strategy to conduct pooled CRISPR-Cas9 screens in mouse primary neurons. Similar to human cell lines, synthetic PR_20_ dipeptides were toxic to mouse primary neurons in a dose-dependent manner (**Fig. 2A**). However, to better model the endogenous production of DPRs, as in patient brains, we also utilized a lentiviral construct expressing codon optimized PR_50_ driven by the neural-specific synapsin promoter (*30*). When expressed in primary neurons, PR_50_ trafficked to the nucleus and elicited neurotoxicity in a time-dependent manner (**Fig. 2 B, C, S3**). We constructed a custom lentiviral sgRNA library to test the top ∼200 genes (5% FDR) found to be genetic modifiers of PR_20_ toxicity in the genome-wide CRISPR screens. The custom library contained ten sgRNAs targeting each mouse gene and ∼1,000 negative control sgRNAs. Using primary neuron cultures isolated from mice constitutively expressing Cas9 (*31*), we used immunoblotting to confirm efficient protein reduction of target genes after transduction with sgRNAs from this library following at least 10 days in culture (**Fig. S4 A-C**).

**Figure 2.**
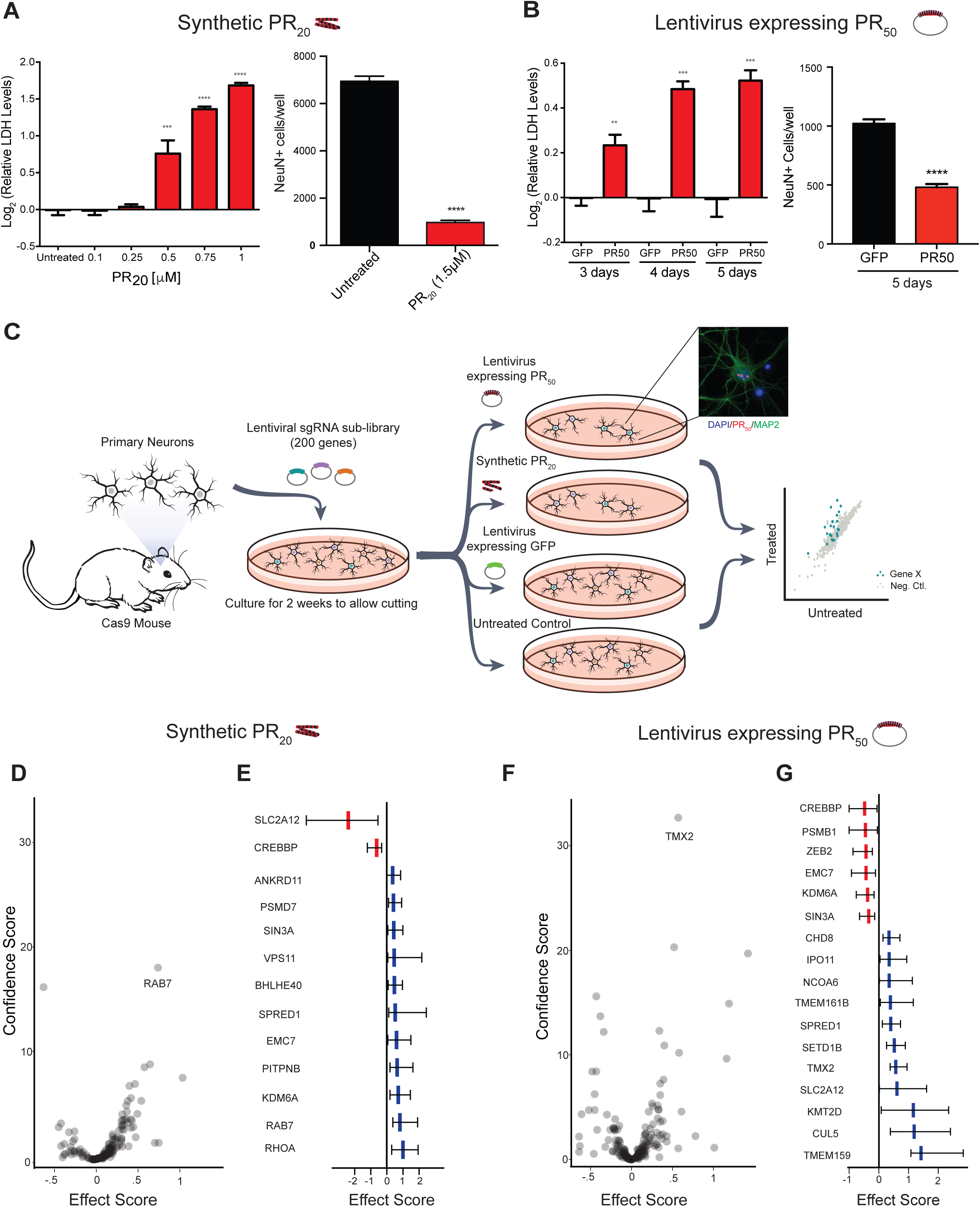
CRISPR-Cas9 knockout screens in primary mouse neurons. **(A)** Synthetic PR_20_ added to primary neuron cultures caused dose-dependent cytotoxicity within 24 hours measured by lactose dehydrogenase (LDH) release or counts of NeuN^+^ neuronal nuclei by immunocytochemistry (two-tailed t-test; **** p < 0.0001, *** p < 0.001, ** p < 0.01). **(B)** Genetically encoded expression of PR_50_ caused time dependent cytotoxicity measured by LDH release or counts of NeuN^+^ neuronal nuclei by immunocytochemistry (two-tailed t-test; **** p < 0.0001, *** p < 0.001, ** p < 0.01). **(C)** CRISPR-Cas9 screening paradigm in primary mouse neuron cultures. Cortical neurons were plated and infected with sgRNA libraries (∼200 genes passing a 5% FDR cutoff from the genome-wide PR screen with 10 sgRNAs/gene + ∼1000 negative control sgRNAs). Four independent screening conditions were conducted, each in replicate: synthetic PR_20_ treated neurons (1.5μM, overnight), untreated neurons, lentiviral expression of PR_50_ (5 days), and lentiviral expression of GFP (5 days) as a control. The abundance of sgRNAs in the surviving cells was measured by sgRNA sequencing. PR50 localized to the nucleus of cultured neurons (blue = DAPI, red = anti-FLAG (PR50), green = anti-MAP2). **(D, F)** Volcano plots representing effect scores and confidence scores of all genes in each screen. **(E, G)** Forest plots of all genes from each screen with a non-zero effect estimate (95% C.I.) with estimated effect in center and error bars representing 95% credible interval of effect estimate. Effect estimate colored in blue if the gene was protective when knocked out and colored in red if it was sensitizing when knocked out.

We performed the primary neuron CRISPR screens two ways – either expressing PR_50_ using lentivirus or treating with exogenous synthetic PR_20_. In this way, we could simultaneously validate hits from the genome-wide screens and disentangle the genes responsible for the uptake and spread of the DPRs throughout the cell from those responsible for mediating the toxic effects once inside the cell. We performed these targeted CRISPR KO screens such that there was a ∼90% reduction in viability after one round of selection; only neurons with the strongest genetic modifiers would become enriched in the selection. We harvested the surviving neurons and assessed sgRNA enrichment compared to the untreated controls (**Fig. 2C**). We identified 13 modifier genes in the synthetic PR_20_ screen and 17 genes in the endogenous PR_50_ expression screen (determined by having a non-zero estimated effect with a 95% credible interval) (**Fig. 2 D-G, Supplementary Table 2).** Strikingly, the genes that validated in this neuronal culture system showed strong specificity to the method of PR delivery to the neurons.

A top hit in the screen for modifiers of synthetic PR_20_ induced toxicity was the endosomal trafficking gene *Rab7a* (**Fig. 2D**). We hypothesize that the endosomal trafficking pathway is necessary for the cellular uptake of the exogenously applied PR_20_. In contrast, none of the trafficking genes present in the mini-library were hits in the screen in which neuronal toxicity was induced by genetically encoded PR_50_ expression. In this screen, the strongest genetic modifiers were genes that encode proteins predominantly localized to the nucleus and ER; a top protective knockout was a poorly characterized ER-resident thioredoxin protein *Tmx2*, which was also a protective genetic modifier in the primary neuron exogenous PR_20_ screen (**Fig. 2F**).

The abundance of strong genetic modifiers involved in ER function and ER stress from our screens suggested that DPR accumulation might induce an ER stress response. To test this hypothesis, we performed RNA sequencing (RNA-seq) on primary neuron cultures transduced with a lentivirus expressing either PR_50_ or GFP (**Fig. 3A**). 126 genes were significantly upregulated and 133 genes significantly downregulated following four days of PR_50_ expression (adjusted p-value < 0.05; **Fig. 3B, S5, Supplemental Table 3**). The most significantly enriched Gene Ontology (GO) category among upregulated genes was ‘apoptotic signaling pathway in response to endoplasmic reticulum stress’ (**Fig. 3C**). These included genes encoding Atf4, Bbc3 (Puma), Chac1, Bax, and the ER Ca^2+^ channel Itpr1, in which loss of function variants have been shown to cause spinocerebellar ataxia 29 (*32*) (**Fig. 3D**). We confirmed a time dependent induction of these ER stress genes by PR_50_ expression using quantitative reverse transcription PCR (qRT-PCR) (**Fig. 3E**). To confirm that these stress responses are conserved in other models of DPR toxicity we performed RNA-seq on primary mouse neurons as well as human K562 cells, each treated with synthetic PR_20_. *ATF4* and related ER stress response genes were induced in each of these models (**Fig. 3F, S6, Supplemental Table 4**). Finally, in addition to observing genetic signatures of ER stress, we also used a pharmacological approach to test the importance of this pathway in response to PR_20_. The small molecule ISRIB is a potent inhibitor of the cellular response to ER stress (*33*) preventing *ATF4* induction by inhibiting eIF2α phosphorylation in human cells under ER-stress (*34*). Pre-treatment of K562 cells with 15nM ISRIB mitigated PR_20_ toxicity (**Fig. 3G**), providing further evidence for a direct role of ER stress in DPR-mediated toxicity.

**Figure 3.**
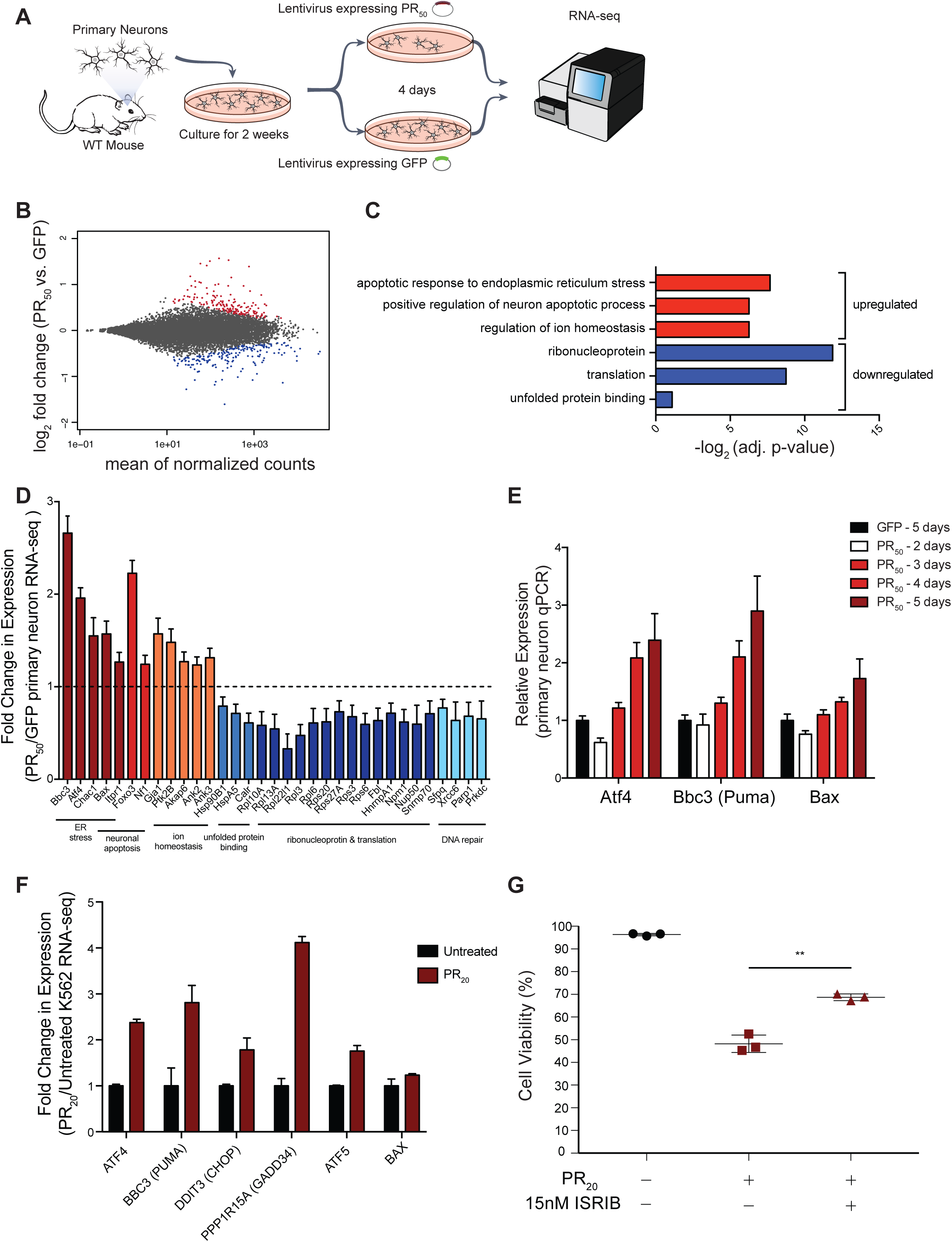
Transcriptional analysis of PR treated primary neurons and K562 cells reveals ER stress signatures. **(A)** RNA sequencing of primary mouse neurons treated with lentiviruses expressing either PR_50_ or GFP under control of the synapsin promoter for 4 days. Differentially expressed genes were determined using DEseq2. **(B)** MA plot showing differential gene expression of PR_50_ expressing neurons compared to control GFP expressing neurons (red = significantly upregulated genes, adjusted p-value < .05; blue = significantly down regulated genes, adjusted p-value < .05). **(C)** Select top GO terms enriched within significantly upregulated genes (red) or significantly downregulated genes (blue). **(D)** Fold change in expression of select genes that were found to be significantly differentially expressed by DEseq2 (adjusted p-value <.001), grouped by GO category. **(E)** qRT-PCR validation of *Atf4*, *Bbc3*, and *Bax* upregulation in PR_50_ expressing neurons compared to GFP expressing neurons. **(F)** RNA sequencing in K562 cells treated for 24 hours with synthetic PR_20_ also show ER stress signatures. Relative expression of significantly differentially expressed genes determined by DESeq2 (adjusted p-value < 0.001) plotted using FPKMs for each gene relative to the untreated population of cells. **(F)** K562 cells pre-incubated with 15nM ISRIB or equal amounts of DMSO were treated with 15μM PR_20_ (red) or untreated (black) for 24 hours, then measured for viability by flow cytometry (FSC/SSC). Each measurement was performed in triplicate with mean and SD plotted by error bars (p-value <.005, two-tailed t-test).

Several lines of evidence have suggested a potential role for ER stress in *C9orf72* pathogenesis. Transcriptomic analyses of *C9orf72* mutant ALS brain tissue reveals an upregulation in genes involved in the unfolded protein response (UPR) and ER stress (*26*). A specific PERK-ATF4 mediated ER stress response was reported in a separate study of c9ALS/FTD patient brain tissue when compared to sporadic ALS patient brain tissue (*35*). Additionally, ER stress and disruptions to ER Ca^2+^ homeostasis were observed in c9ALS/FTD patient derived iPSC motor neurons (*36*). Because *C9orf72* mutations could cause disease pathology by several different mechanisms, including loss of function, RNA toxicity, and/or protein toxicity from the DPRs (*37*), it has remained unclear which molecular drivers induce the ER stress response. The ATF4-driven ER stress response we observed in primary neuron cultures expressing PR_50_ and the enrichment of genes functioning at the ER as hits in our CRISPR screens suggest that this stress response may be a major driver of pathogenesis and, importantly, is likely driven by the production of arginine-rich DPRs instead of by loss of C9orf72 function or toxicity from repeat-containing RNAs.

Given that the main transcriptional stress response to endogenous and exogenous DPR treatment was the ER stress response, we chose to further investigate *TMX2*, a gene encoding a poorly characterized ER-resident protein that was one of the strongest protective hits in both the PR and GR genome-wide CRISPR screens in K562 cells and in the targeted primary neuron screen expressing PR_50_. We first confirmed that the sgRNAs targeting *TMX2* effectively knocked out the gene in K562 cells and primary neurons cultured from the Cas9 mouse (**Fig. 4A,C**). Strikingly, reduction of *TMX2* levels was sufficient to markedly suppress *C9orf72* DPR toxicity in both K562 cells (**Fig. 4B**) and primary mouse neurons (**Fig. 4D**).

**Figure 4.**
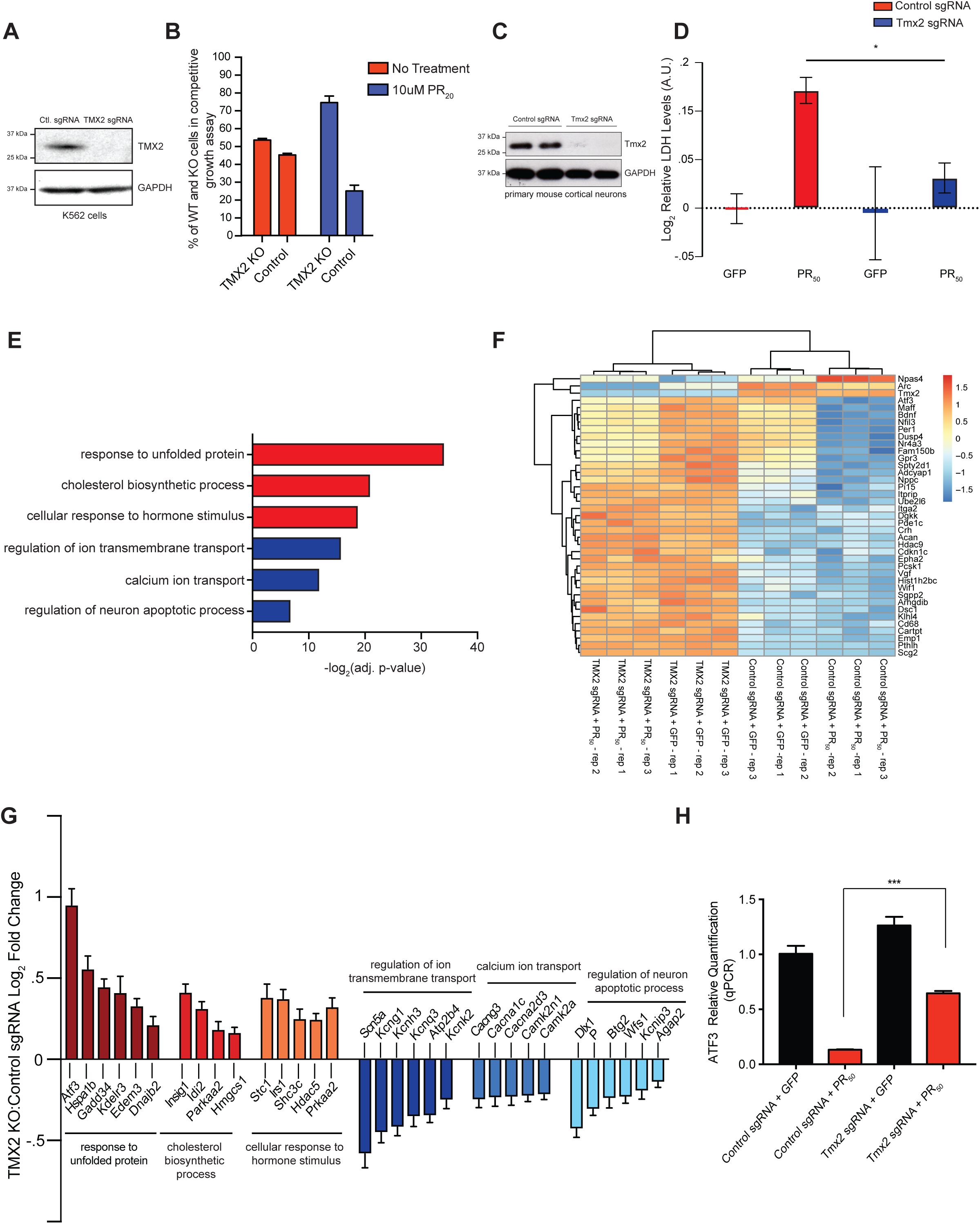
*TMX2* reduction is protective against PR mediated toxicity. **(A)** *TMX2* KO in K562 cells confirmed by immunoblot. **(B)** Enrichment of *TMX2* KO cells compared to WT in a competitive growth assay after synthetic PR_20_ treatment. Cells measured 48 h after PR_20_ treatment by flow cytometry. **(C)** *Tmx2* KO confirmed in primary mouse neurons by immunoblot. **(D)** *Tmx2* KO protected primary neurons against lentiviral expression of PR_50_, measured by LDH release after 4 days of PR_50_ expression. **(E)** Select GO term enrichment (red = upregulated genes, blue = downregulated genes) of differentially expressed genes (adjusted p-value <.001) determined by comparing RNA-seq results between control and *Tmx2* KO neuron cultures expressing PR_50_. **(F)**. Differential gene expression was determined using DEseq2, and the top differentially expressed genes as ranked by adjusted p-value were clustered using R package pheatmap. **(G)** Log_2_ fold change of differentially expressed genes (adjusted p-value <.001) enriched in select GO categories **(H)** qRT-PCR validation of *Atf3* expression changes after PR_50_ expression in control and *Tmx2* KO neurons.

TMX2 is a transmembrane protein that localizes to the mitochondrial-associated membrane (MAM) of the ER (*38*). It is a member of the protein disulfide isomerase (PDI) family, but it is unclear whether it is catalytically active since it harbors an SXXC motif instead of the canonical CXXC motif thought to be necessary to reduce disulfide bonds (*39*). To define the mechanism by which *TMX2* reduction mitigates DPR toxicity we performed RNA-seq on primary Cas9^+^ neurons infected with control sgRNAs or sgRNAs targeting *TMX2,* with or without PR_50_ expression (**Fig. 4E-G, S7**). Upon PR_50_ expression, *Tmx2* KO neurons upregulated a distinct set of pro-survival unfolded protein response (UPR) pathway genes and downregulated calcium binding and apoptotic genes, compared to primary neurons expressing control sgRNAs (**Fig. 4G**). We validated by qRT-PCR one of the upregulated UPR genes in the *Tmx2* KO neurons, *Atf3*, which has been previously shown to be protective upon overexpression in mouse models of ALS (*40*) and neuronal models of excitotoxicity (*41*) (**Fig. 4H**). These results suggest that *TMX2* KO may be protective by modulating the ER stress response elicited by the DPRs.

Here, we used comprehensive CRISPR-Cas9 screens in human cells with further validation screens in primary neurons to discover modifiers of *C9orf72* DPR toxicity. We not only identified nucleocytoplasmic transport machinery, confirming previous studies in model organisms, but also identified new genes that point to ER stress as critical for c9FTD/ALS pathogenesis. The gene knockouts that mitigated toxicity, such as *TMX2*, could represent therapeutic targets. It is notable that *TMX2* KO was able to confer near complete protection from PR toxicity. More generally, we anticipate that these types of CRISPR screens in human cells and primary neurons will be a powerful addition to the experimental toolbox to study mechanisms of neurodegenerative disease.

## Acknowledgments

We thank Emily Crane and Billy Li for help with RNA-seq and helpful discussions, Kyuho Han for helpful discussions and image analysis. This work was supported by NIH grants R35NS097263 (A.D.G.) and DP2HD084069 (M.B.), the National Science Foundation Graduate Research Fellowship (N.J.K.), The National Human Genome Research Institute Training Grant (M.S.H.), the Robert Packard Center for ALS Research at Johns Hopkins (A.D.G.), Target ALS (M.B., A.D.G.), and the Stanford Brain Rejuvenation Project of the Stanford Neurosciences Institute (M.B., A.D.G.). RNA sequencing data will be deposited with NCBI Geo upon publication.

## Supplementary Materials

CRISPR-Cas9 screens in human cells and primary neurons identify modifiers of *C9orf72* dipeptide repeat protein toxicity

Michael S. Haney^1^*, Nicholas J. Kramer^1,2,^*, David W. Morgens^1^, Ana Jovičić^1,3^, Julien Couthouis^1^, Amy Li^1^, James Ousey^1^, Rosanna Ma^1^, Gregor Bieri^1,2^, Michael C. Bassik^1,#^, Aaron D. Gitler^1, #^

## Supplementary Methods

### Cell Culture

K562 cells (ATCC) were cultured in RPMI-1640 (Gibco) media with 10% fetal bovine serum (FBS) (Hyclone), penicillin (10,000 I.U./mL), streptomycin (10,000 μg/mL), and L-glutamine (2 mM). Cells were grown in log phase by maintaining the population at a concentration of 500,000 cells per mL. K562 cells were maintained in a controlled humidified incubator at 37°C, with 5% CO_2_.

Primary mouse neurons were dissociated into single cell suspensions from E16.5 mouse cortices using a papain dissociation system (Worthington Biochemical Corporation). Neurons were seeded onto poly-L-lysine coated plates (0.1% w/v) and grown in Neurobasal media (Gibco) supplemented with B-27 serum-free supplement (Gibco), GlutaMAX, and Penicillin-Streptomycin (Gibco) in a humidified incubator at 37°C, with 5% CO_2_. Half media changes were performed every 4-5 days, or as required.

### Lentivirus production

HEK293T cells were cultured under standard conditions (DMEM + 10% FBS + Penicillin-Streptomycin.) and used to package lentiviral particles following standard protocols with 3rd generation packaging plasmids. Lentiviral containing media was harvested after 48hrs., centrifuged at 300g for 5 min. to remove cellular debris, and concentrated 10-fold using Lenti-X concentrator (Clontech) before adding to cell cultures.

### Immunocytochemistry and antibodies

Cells were grown on poly-L-lysine coated glass coverslips (0.1% w/v) in standard multiwell cell culture plates and were stained using standard immunocytochemistry techniques. Briefly, cells were fixed with 4% formaldehyde, permeabilized with 0.1% Triton X-100, blocked with 5% normal goat serum, and stained with the following antibodies: mouse monoclonal anti-FLAG (1:1000, M2 Sigma cat.# F1804), mouse monoclonal anti-MAP2A (1:1000, Millipore cat.# MAB378), or mouse monoclonal anti-NeuN (1:1000, Millipore cat.# MAB377). Coverslips were mounted using Prolong Diamond Antifade Mountant with DAPI (Thermo Fisher Scientific). Images were acquired using either a Leica DMI6000B inverted fluorescence microscope with a 40X oil immersion objective or a confocal Zeiss LSM710 microscope with a 63X oil immersion objective.

### Genome-wide CRISPR-Cas9 screens in K562 cells

The 10-sgRNA-per-gene CRISPR/Cas9 deletion library was synthesized, cloned, and infected into Cas9 expressing K-562 cells as previously described (Morgens et al., 2017). Briefly ∼300 million K562 cells stably expressing SFFV-Cas9-BFP were infected with the 10 guide/gene genome-wide sgRNA library at an MOI < 1. Infected cells underwent puromycin selection (1ug/mL) for 5 days after which point puromycin was removed and cells were resuspended in normal growth media without puromycin. After selection sgRNA infection was measured as >90% of cells as indicated measuring mCherry positive cells by flow-cytometry. Sufficient sgRNA library representation was confirmed by deep sequencing after selection. Cells were maintained for three weeks at 1000x coverage (∼1000 cells containing each sgRNA) at a concentration of 500,000 cells/mL for the duration of each screen. Cells were split into two conditions: one control group that remained untreated, and one group that was treated with the respective synthetic DPR (PR_20_ at 8μM or GR_20_ at 10μM). This treatment occurred four times over three weeks, allowing for cells to recover to ∼90% viability after each round of treatment. At the end of each screen genomic DNA was extracted for all screen populations separately according to the protocol included with QIAGEN Blood Maxi Kit and deep sequencing on an Illumina Nextseq was used to monitor library composition. Guide composition was sequenced and compared to the plasmid library using casTLE(Morgens et al., 2016) version 1.0 available at https://bitbucket.org/dmorgens/castle.

### K562 competitive growth assays

K562 cells stably expressing Cas9 were infected with sgRNAs targeting each gene of interest for validation from the genome wide screen. sgRNA plasmids also encoded for GFP expression. Cells were selected for stable sgRNA expression using puromycin (1 ug/mL) and were confirmed to be GFP positive before use in competitive growth assays. Equal numbers of GFP^+^/sgRNA expressing cells were co-cultured with wildtype/GFP-cells, and were subsequently treated with 8 μM PR_20_. The percentage of GFP+ cells were quantified by fluorescence activated cell sorting using a BD Accuri C6 flow cytometer after 24 hours.

### Pre-treatment of ISRIB in PR_*20*_ exposed K562s

ISRIB (SIGMA SML0843) was dissolved in DMSO and added to the K562 cell line at a concentration of 15nM 3 hours before treatment with PR_20_. An equal amount of DMSO was added to cells that did not get pre-treated with ISRIB. 15uM of PR_20_ was then added to these cells and a third group of K562s were not treated with ISRIB or PR_20_ to serve as an untreated control. 24 hours after PR_20_ treatment cells were measured using the BD Accuri C6 flow cytometer. All experiments were performed in triplicate.

### CRISPR-Cas9 screen for DPR toxicity in primary mouse cortical neurons

Cortical tissue from mouse E.16.5 embryos [*Gt(ROSA)26Sor*^*tm1.1(CAG-cas9*,-EGFP)Fezh*^/J; Jackson Laboratory Stock No: 024858] was dissected and dissociated into a single cell suspension using a papain dissociation system (Worthington Biochemical Corporation). 7 million cells were seeded onto 10cm dishes respectively in Neurobasal media (Gibco) supplemented with B-27 serum-free supplement (Gibco), GlutaMAX (Gibco), and Penicillin-Streptomycin (Gibco). Half the culture media was replenished every 4-5 days. One day after seeding, neurons were infected with a lentiviral library containing ∼3000 sgRNA elements consisting of 178 genes (5% FDR for PR_20_ screen hits), 10 sgRNAs/gene, and 2000 negative control sgRNAs, synthesized and cloned as previously described (Morgens et al., 2016). Using a titer resulting in a 50-70% infection rate. Cas9-mediated genome editing was allowed to occur for 2 weeks in culture, after which PR_20_ synthetic dipeptides or PR_50_ expressing lentivirus were applied. Synthetic DPR screen: half the cells were left untreated (control group), and the other half were treated with 1.5 μM synthetic PR_20_ for 24hr, which resulted in approximately 90% cell death. Lentiviral mediated DPR screen: half the cells were infected with a control lentivirus expressing eGFP from the synapsin promoter (control group), and the other half were infected with a lentivirus expressing codon optimized PR_50_ from the synapsin promoter. Lentiviral PR_50_ resulted in approximately 90% cell death between 4-6 days. To harvest DNA from the remaining live cells, cultures were washed 3 times in PBS, then treated with DNaseI (Worthington Biochemical Corporation) for 10 minutes at 37° C (to remove any remaining gDNA from dead neurons), and then washed again 3 times in PBS to remove residual DNaseI. Genomic DNA was harvested using a DNeasy Blood and Tissue Kit (QIAGEN) including Proteinase K digestion to inactivate residual DNaseI. sgRNA amplification and library preparation for deep sequencing was performed as described above.

### RNA-sequencing

Total RNA quality control was performed using the Eukaryote Total RNA Nano assay on the Agilent 2100 Bioanalyzer System for all RNA samples prior to library preparation for RNA sequencing. mRNA libraries were prepared for Illumina paired-end sequencing using the Agilent SureSelect Strand Specific RNA-Seq Library Preparation kit on an Agilent Bravo Automated Liquid Handling Platform. Libraries were sequenced on an Illumina NextSeq sequencer. Alignment of RNA-sequencing reads to the transcriptome was performed using STAR with ENCODE standard options, read counts were generated using rsem, and differential expression analysis was performed in R using DEseq2 package (*42*). All bioinformatics analyses were performed on Sherlock, a Stanford HPC cluster.

### qRT-PCR

cDNA was reverse transcribed from total RNA samples using the High Capacity cDNA Reverse Transcription Kit (ThermoFisher Scientific). Taqman assays (Applied Biosystems) for ATF3, ATF4, BBC3, and BAX were used in qPCR reactions using Taqman Universal PCR Master Mix (ThermoFisher Scientific) and StepOne or QuantStudio 3 real time PCR machines.

### Western Blotting

Protein lysates were prepared using RIPA buffer supplemented with 1X Halt Protease Inhibitor Cocktail. Crude lysates were centrifuged at 12,500xg for 10 min. at 4°C to remove cellular debris. Clarified lysates were quantified using Pierce BCA protein assay. Equal amounts of protein were subjected to SDS-PAGE, transferred to nitrocellulose membranes, and immunoblotted following standard protocols. The following antibodies were used: mouse anti-GAPDH (1:5000, clone GAPDH-71.1, Sigma G8795), rabbit polyclonal anti-TMX2 (1:1000, Novus NBP1-87305).

### Cytotoxicity Assays

K562 cell viability after DPR treatment was measured by flow cytometry using a BD Accuri C6 flow cytometer by measuring number of events in forward scatter and side scatter gates. Cytotoxicity in primary neuron cultures was measured both by lactose dehydrogenase (LDH) release assays (Promega, CytoTox 96^®^ Non-Radioactive Cytotoxicity Assay) and by NeuN+ quantification of surviving neurons using immunocytochemistry. To quantify numbers of NeuN+ neurons, cells were fixed and stained as described above, and quantified using ImageJ. Briefly, thresholds were applied to image stacks in order to detect NeuN positive neuronal nuclei, which were then automatically counted using built in ImageJ plugins.

## Supplementary Figures

**Figure S1.**
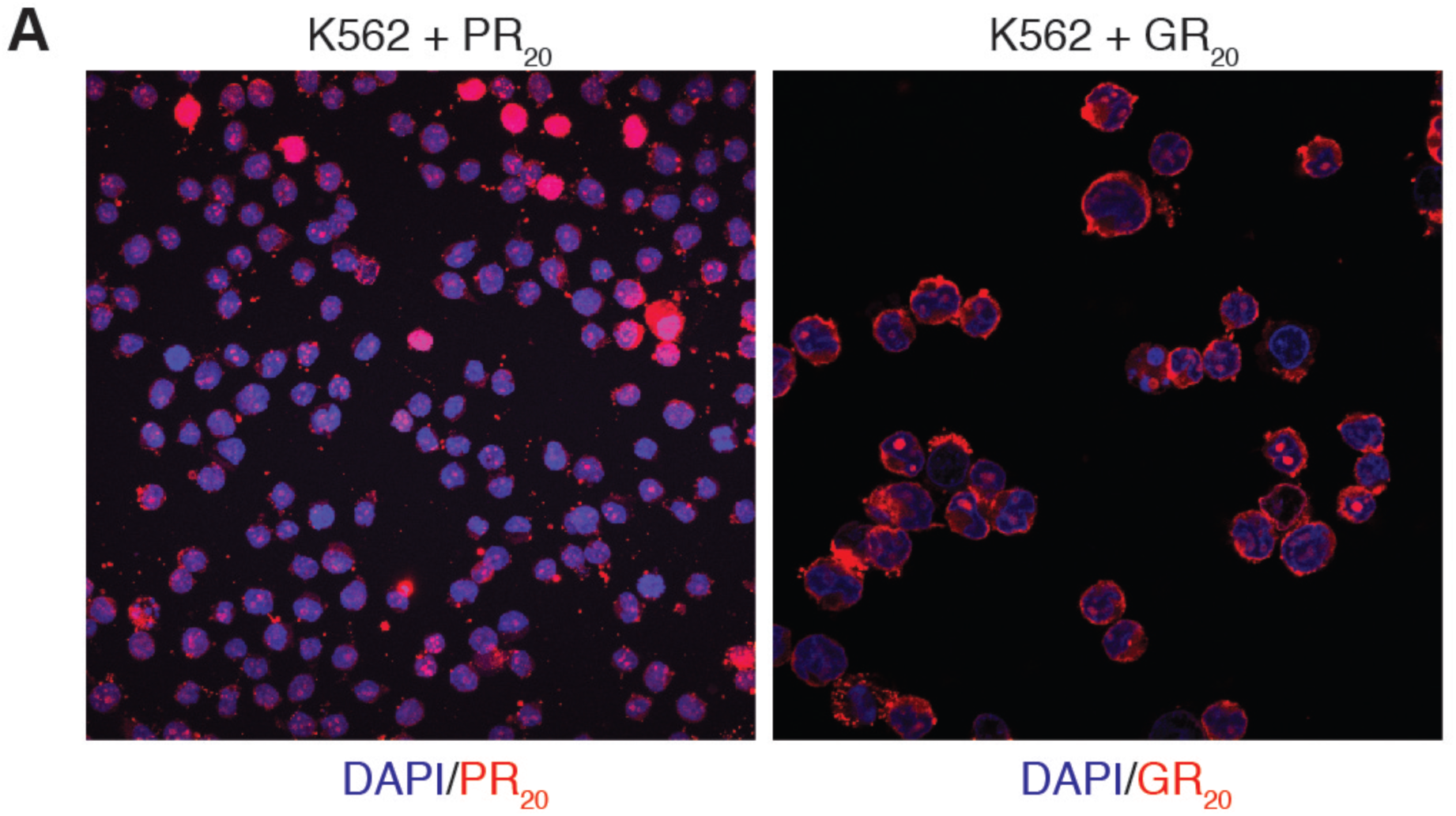
Localization of FLAG-tagged DPRs applied exogenously to K562 cells. PR_20_ and GR_20_ localized to the nucleus of K562 cells as visualized by immunocytochemistry (blue = DAPI, red = anti-FLAG).

**Figure S2.**
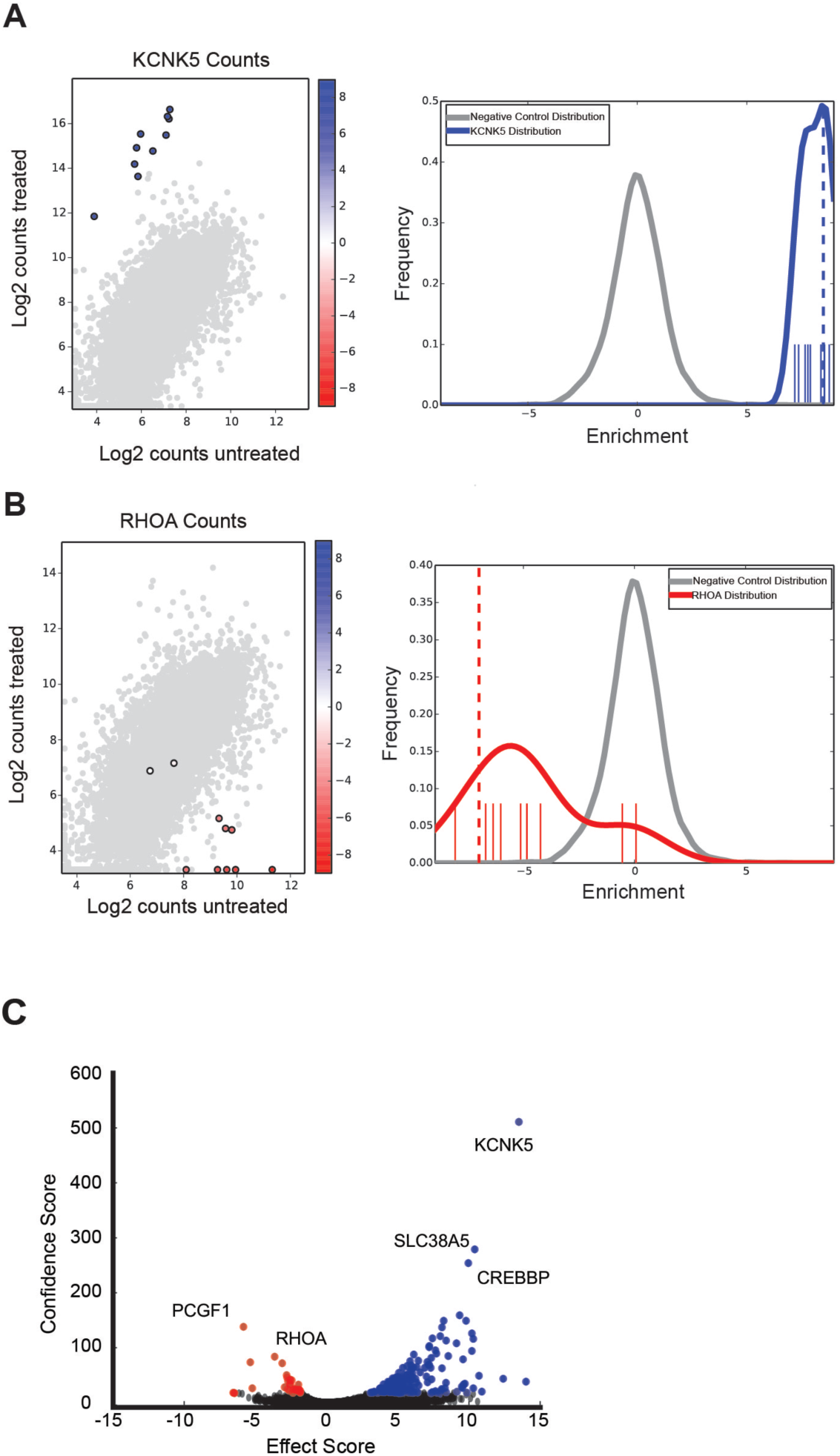
Distributions of sgRNAs counts from an example protective genetic modifier (*KCNK5*) and sensitizing genetic modifier (*RHOA*). **(A)** Distribution of sequencing from 10 *KCNK5* sgRNAs (blue) compared to distribution of sequencing counts of all negative control sgRNA (grey) from PR_20_ screen replicate 1. **(B)** Distribution of sequencing from 10 *RHOA* sgRNAs (red) compared to distribution of sequencing counts of all negative control sgRNA (grey) from PR_20_ screen replicate 1. Using CasTLE these sgRNA count distributions were used to generate effect scores, confidence scores and p-values for each gene **(see Supplemental Table 1)**. **(C)** Volcano plot of effect vs. confidence scores for all human genes in the PR_20_ screen. Colored in blue are all the genes conferring resistance to PR_20_ when knocked out (10%FDR) and colored in red are all the genes conferring sensitivity to PR_20_ when knocked out (10% FDR).

**Figure S3.**
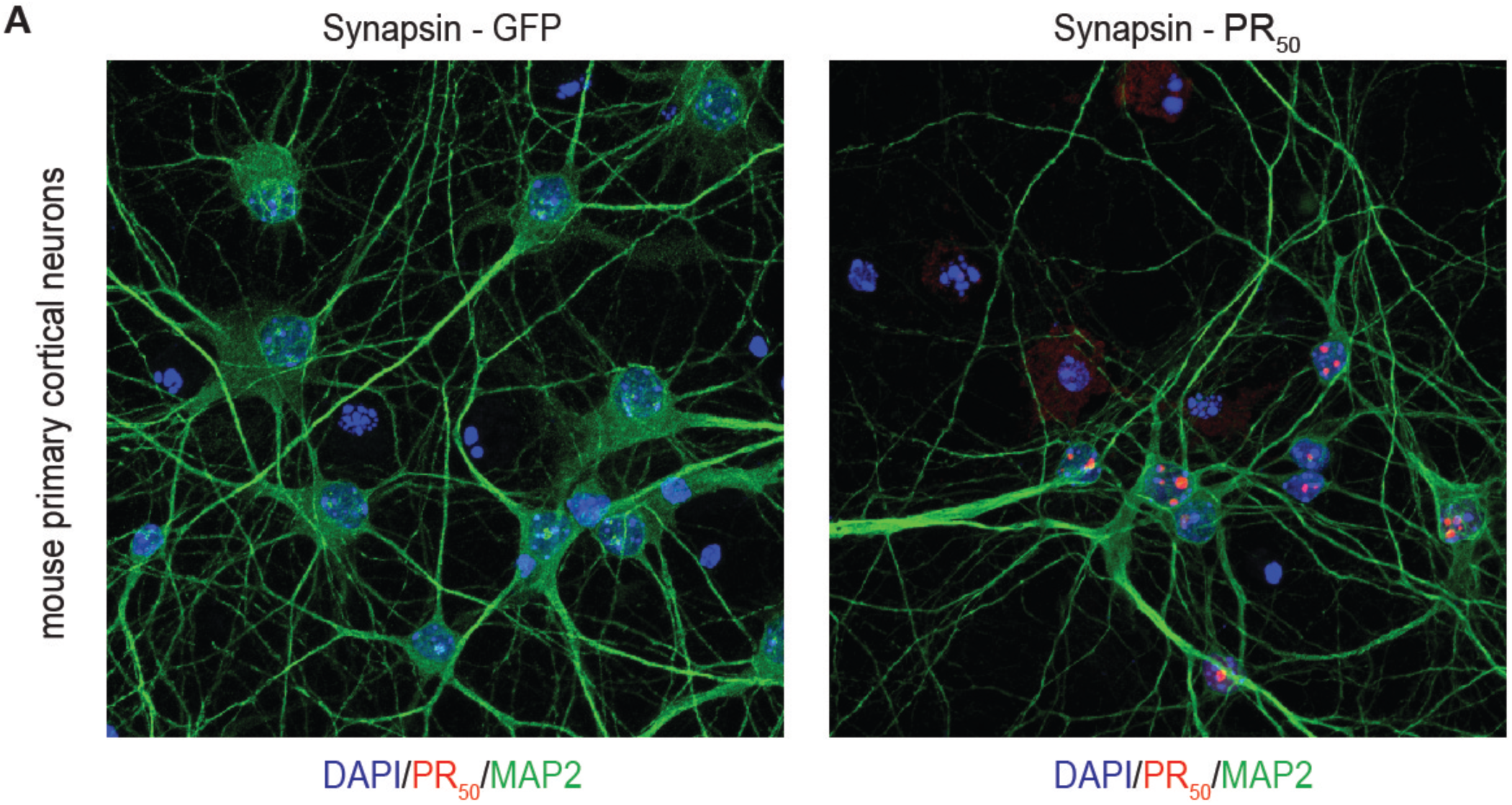
PR_50_ localization in primary mouse neurons. Lentiviral expression of GFP or PR_50_-FLAG using a synapsin promoter in primary neurons. Immunocytochemistry was used to visualize PR_50_ localization (blue = DAPI, red = anti-FLAG, green = anti-MAP2).

**Figure S4.**
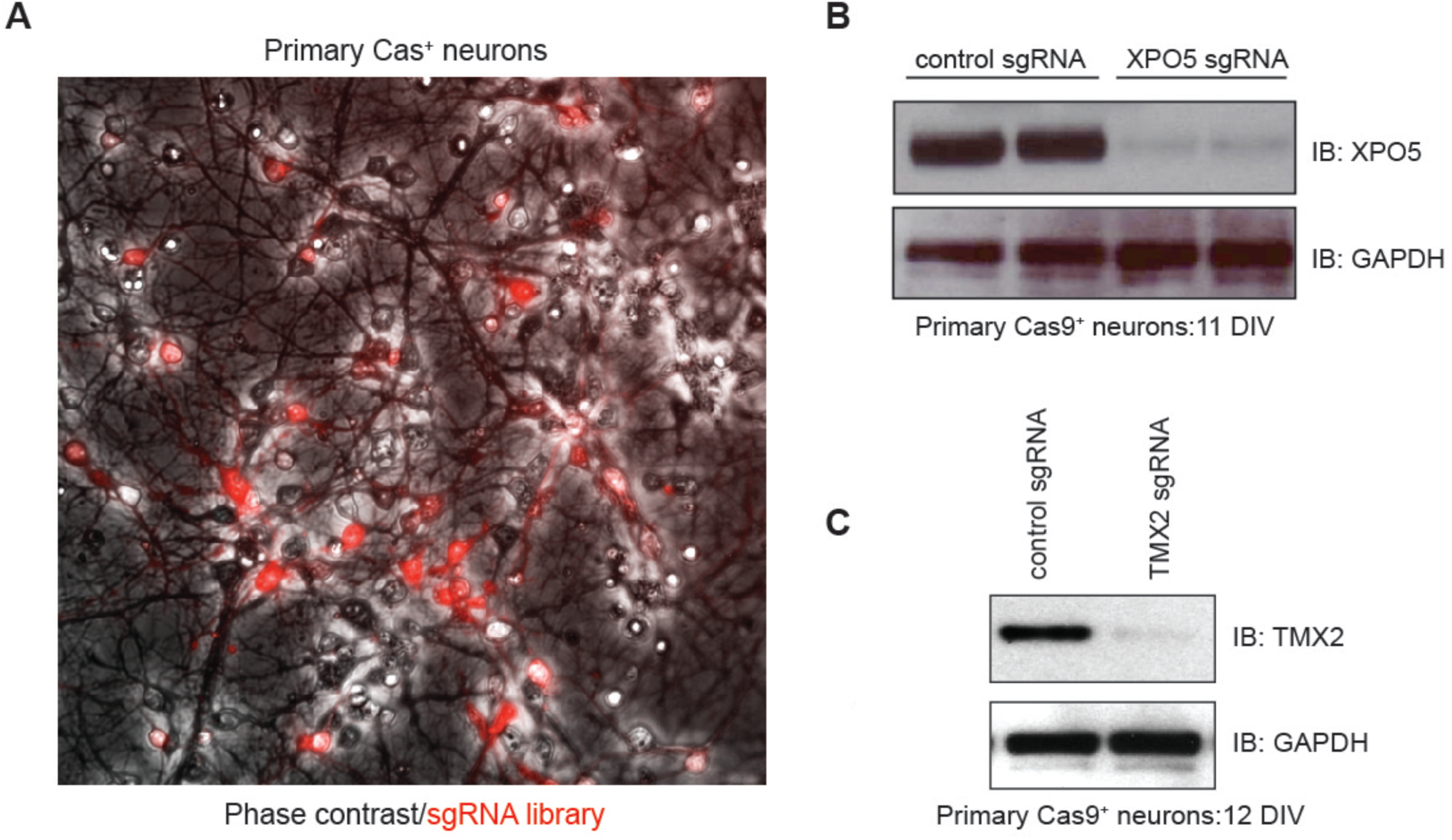
Custom mouse sgRNA library infection and gene KO in primary neurons. **(A)** Visualization of the 3000 sgRNA lentiviral library expressing mCherry in infected primary mouse neurons (grey = phase contrast, red = mCherry; live cells). **(B, C)** Validation of target protein reduction in Cas^+^ primary neurons using sgRNAs targeting *Xpo5* and *Tmx2*. Reduced abundance of target protein in primary neurons as measured by western blot was observed after more than 10 days of sgRNA expression.

**Figure S5.**
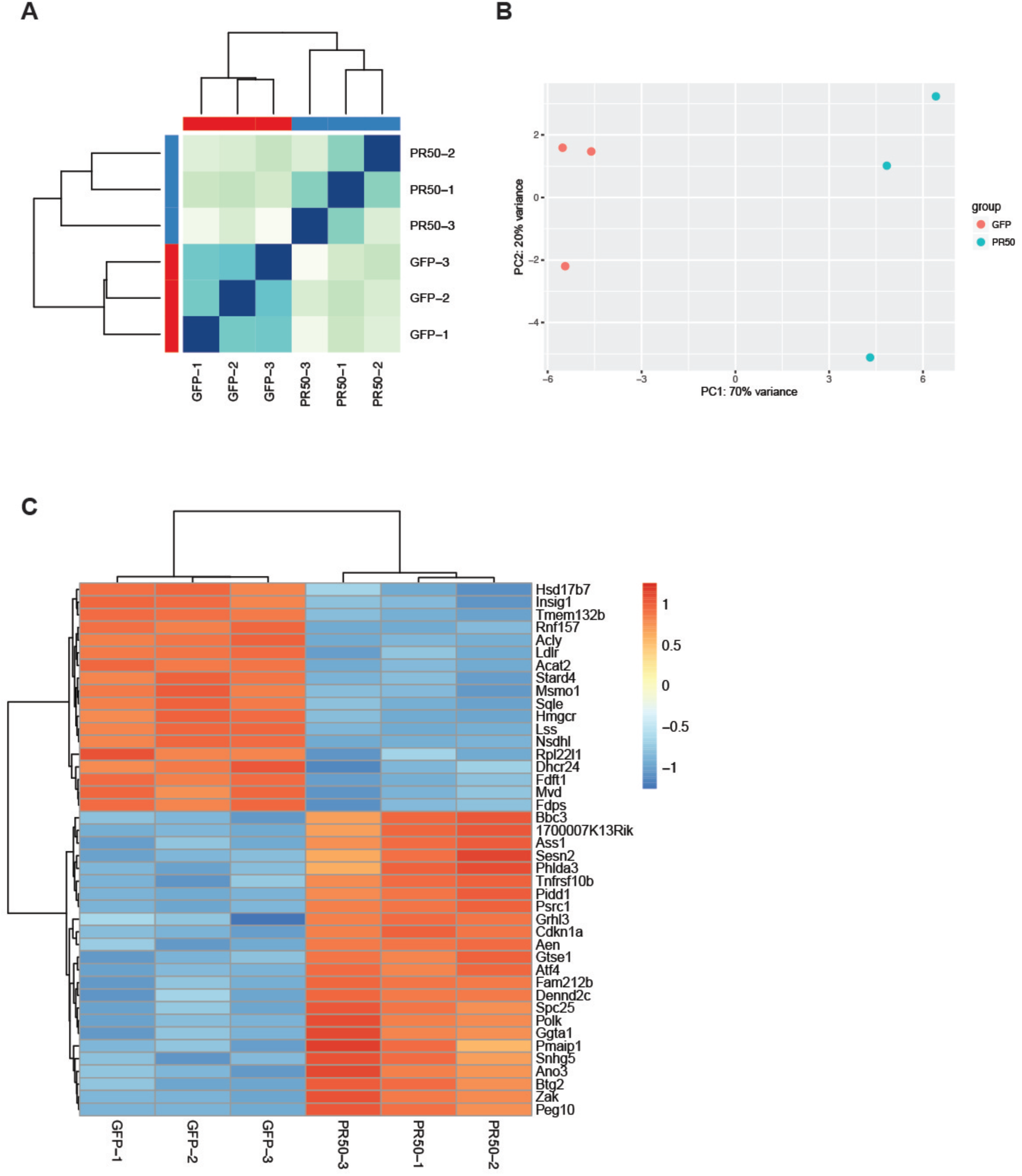
Quality control metrics of RNA-seq data from WT neurons expressing GFP and PR_50_. **(A)** Sample distance matrix using hierarchical clustering with Euclidian distance metric of total transcriptome of each sample. **(B)** Principal component analysis of total transcriptome of each sample. **(C)** Clustered heat map showing normalized expression levels of top 40 genes sorted by adjusted p-value. All analyses were performed using DEseq2 in R **(Supplemental Table 3)**.

**Figure S6.**
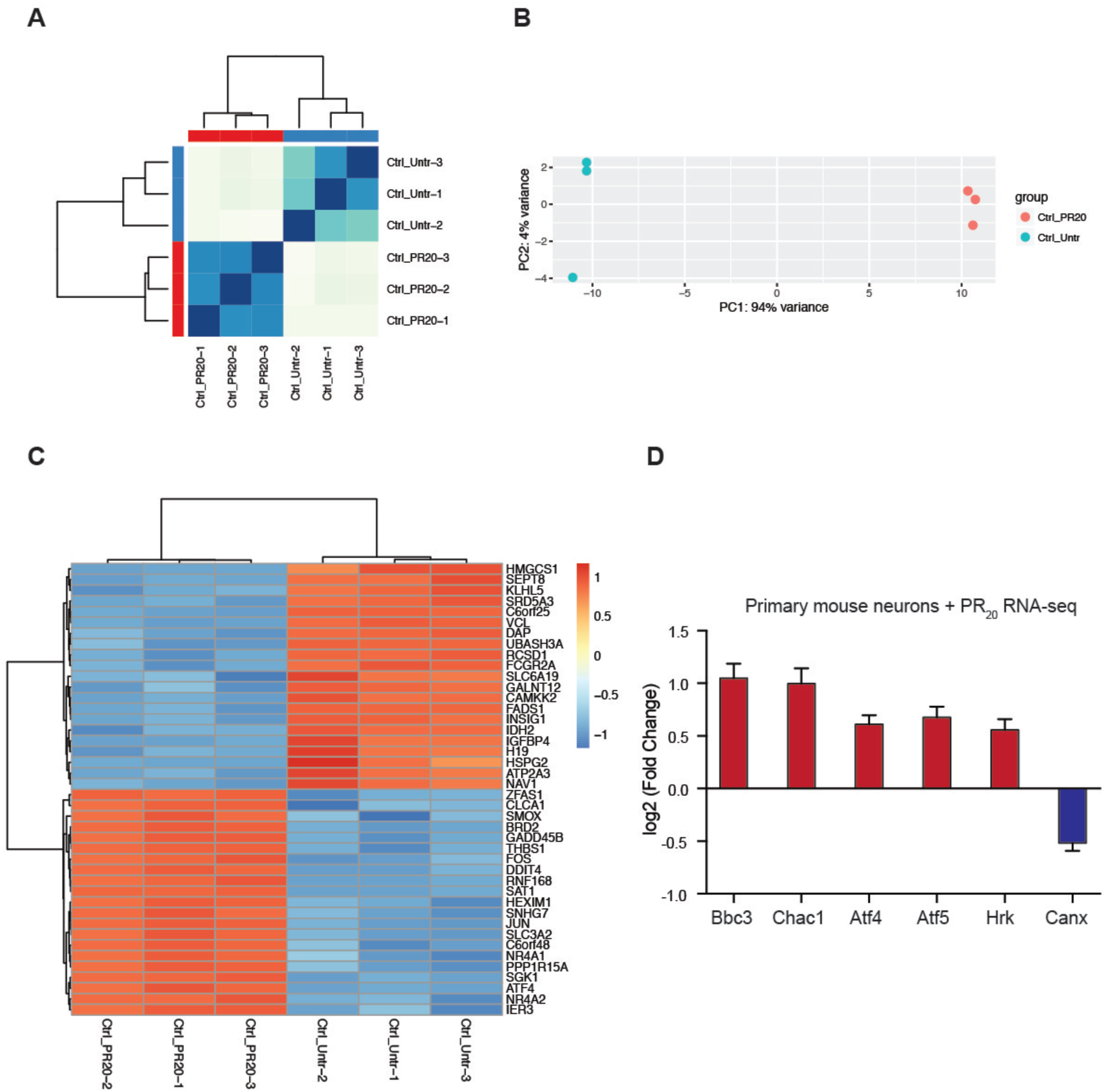
Quality control metrics of RNA-seq data from K562 cells treated with 10 μM PR_20_ and untreated cells. **(A)** Sample distance matrix using hierarchical clustering with Euclidian distance metric of total transcriptome of each sample. **(B)** Principal component analysis of total transcriptome of each sample. **(C)** Clustered heat map showing normalized expression levels of top 40 genes sorted by adjusted p-value. All analyses were performed using DEseq2 in R **(Supplemental Table 4)**. (D) Fold change of select ER-stress related, differentially expressed genes determined by DEseq2 (adjusted p-value <0.001) from RNA-seq data from primary mouse neurons treated overnight with 1.5μM PR_20_ **(Supplemental Table 5)**.

**Figure S7.**
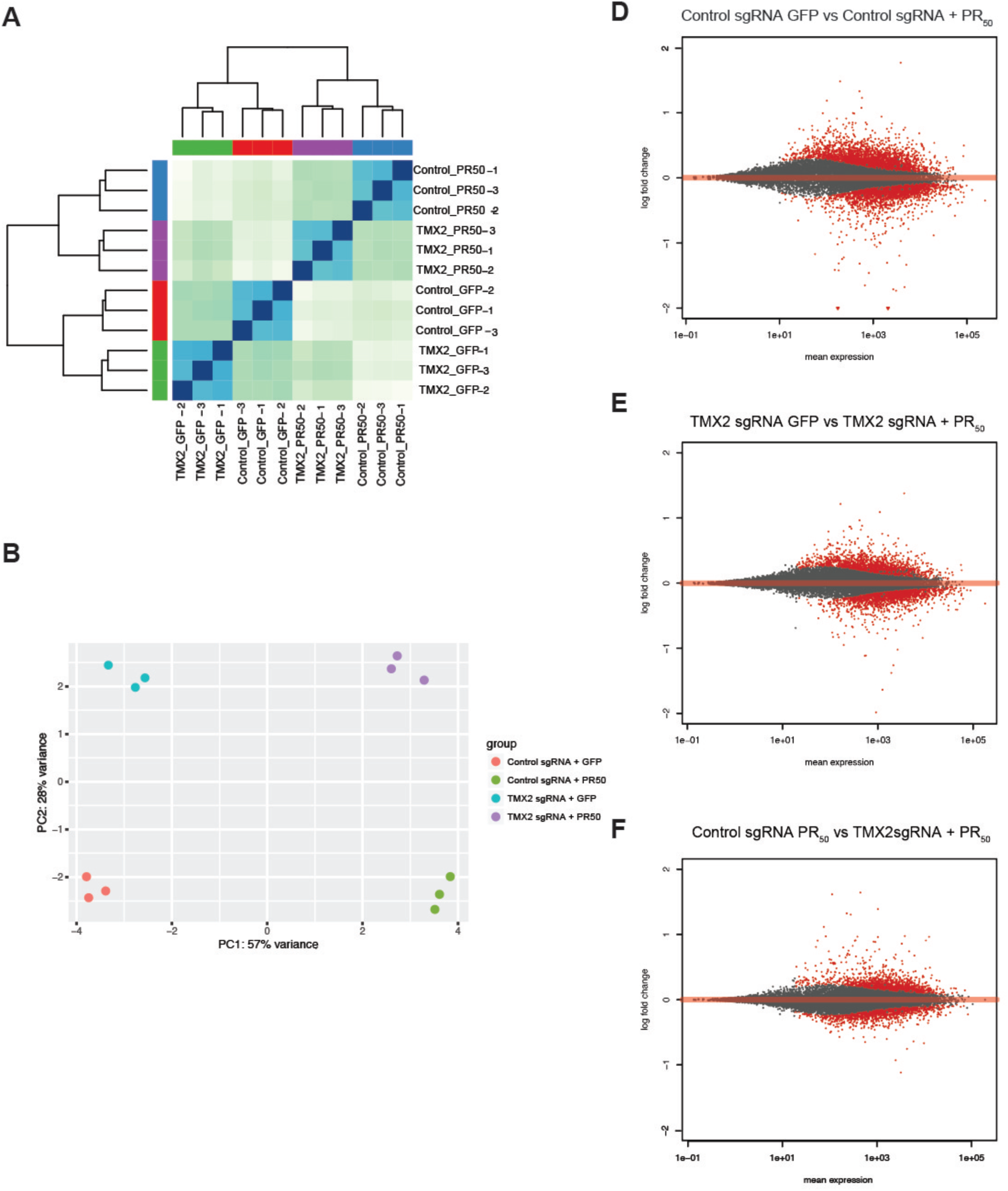
Quality control metrics of RNA-seq data from *Tmx2* KO neurons expressing PR_50_ or GFP. **(A)** Sample distance matrix using hierarchical clustering with Euclidian distance metric of total transcriptome of each sample. **(B)** Principal component analysis of total transcriptome of each sample. MA plot showing abundance of differentially expressed genes in red (adjusted p-value <0 .05) for pairwise comparisons of **(C)** control sgRNA neurons expressing PR_50_, and neurons expressing GFP, **(D)** *Tmx2* KO neurons expressing PR_50_ and *Tmx2* KO neurons expressing GFP, and **(E)** *Tmx2* KO neurons expressing PR_50_ and control sgRNA neurons expressing GFP **(Supplemental Table 7)**.

